# Hypoxia-induced regulation of zDHHC23 opens avenues for new biomarkers for NON MYCN-amplified neuroblastoma

**DOI:** 10.1101/2025.11.03.686197

**Authors:** Sally O. Oswald, Leonard A. Daly, Kim Clarke, Philip J. Brownridge, Barry Pizer, Ian A. Prior, Violaine Sée, Claire E. Eyers

## Abstract

Neuroblastoma is a highly metastatic paediatric malignancy with poor prognosis, and remains a leading cause of paediatric cancer mortality. Current risk stratification is disproportionately reliant on *MYCN* amplification, a feature present in only ∼25% of cases. This narrow focus neglects the majority of patients that have non-*MYCN* amplified tumours, limiting opportunities for therapeutic innovation. To define molecular drivers of metastasis in non*-MYCN* amplified neuroblastoma and identify putative prognostic markers, we leveraged our validated *in vivo* chick embryo xenograft model to profile oxygen-sensitive transcriptional changes, revealing a cohort of ∼400 genes associated with metastatic competence. Integrative survival analysis using two independent patient cohorts identified 59 genes predictive of event-free survival in non-*MYCN* amplified disease. Among these, the poorly characterized Zinc Finger DHHC-Type Palmitoyltransferase 23 (zDHHC23) emerged as a robust prognostic candidate.

Proteomic interrogation of zDHHC23 under varying oxygen conditions uncovered dynamic interactome remodelling, notably hypoxia-attenuated association with the 26S proteasome and the TIM23 mitochondrial import complex. These findings suggest zDHHC23 as a hypoxia-responsive regulator linked to metastatic signalling, and a promising target for biomarker development and therapeutic intervention in high-risk, non-MYCN amplified neuroblastoma.

## Introduction

Neuroblastoma (NB) is the most common malignant solid tumor diagnosed in children, accounting for ∼15% of all childhood cancer-related deaths. Its clinical trajectory is strikingly heterogeneous, ranging from spontaneous regression to aggressive metastatic dissemination refractory to multimodal therapy (1, 2). Despite decades of clinical refinement, current treatment regimens - anchored in chemotherapy, surgery, radiotherapy, and immunotherapy - remain largely non-specific and frequently toxic, imposing a substantial burden of long-term morbidity on survivors whose organs are still developing (3-6). Adverse effects such as sensorineural hearing loss, hypothyroidism, epilepsy, and renal impairment are common (7, 8), underscoring the urgent need for more targeted and less deleterious therapeutic strategies (9).

Since 2008, the International Neuroblastoma Risk Group (INRG) classification system has provided a framework for stratifying patients based on seven clinical and genomic parameters, including *MYCN* amplification, tumour stage, and chromosomal aberrations (10). However, this system remains imperfect. *MYCN* amplification, while a cornerstone of current risk assessment, is present in only ∼25% of cases and fails to capture the molecular complexity of the remaining majority (11-14). Consequently, disease severity is frequently underestimated, particularly in non-*MYCN* amplified tumours, where the molecular drivers of metastasis remain poorly defined. This gap is clinically consequential: over half of NB patients present with metastatic disease at diagnosis (most commonly to bone, bone marrow, liver, and CNS), and relapse with distant metastases is associated with survival rates below 5% (15). A more accurate risk estimation of individual patients at diagnosis is therefore essential for appropriate treatment stratification and for harm minimisation.

To address this crucial gap in the ability to predict outcome and therapeutic response, we investigated the molecular determinants of metastasis in non-*MYCN* amplified NB using our previously validated chick embryo chorioallantoic membrane (CAM) xenograft model (16). The CAM offers a highly vascularised, immunodeficient microenvironment conducive to rapid tumour engraftment and haematogenous dissemination, enabling real-time interrogation of metastatic potential. Compared to rodent models, the chick embryo system is cost-effective, ethically compliant with the 3Rs, and permits high-throughput *in vivo* screening. We have previously demonstrated that hypoxic preconditioning of NB cells induces metastatic competence in this model (16).

Building on this foundation, we performed transcriptomic profiling of SK-N-AS-derived primary NB tumours preconditioned under hypoxia (1% O_2_) versus normoxia (21% O2), identifying candidate genes associated with metastatic behaviour. Of particular interest, we observed upregulation of *Zinc Finger DHHC-Type Palmitoyltransferase 23* (*zDHHC23*) in metastatic tumors, revealing it to be a good predictor of patient survival in non-*MYCN* amplified NB patients. zDHHC23 belongs to the DHHC family of palmitoyltransferases, which catalyse reversible S-palmitoylation, a lipid modification that governs protein localisation, trafficking, and stability (17). Despite its putative regulatory role, zDHHC23 remains poorly characterised, with only one known substrate (KCNMA1) reported to date (18). To elucidate the functional relevance of zDHHC23 in NB, we interrogated its hypoxia-modulated interactome, post-translational modification (PTM) landscape, and subcellular localisation. Our findings provide novel insight into zDHHC23-mediated signalling and suggest this enzyme as a promising candidate for biomarker development and therapeutic targeting in high-risk, non-MYCN amplified neuroblastoma.

## Materials and Methods

### Reagents

Tissue culture reagents were purchased from Gibco (Thermo Fisher Scientific). Powdered chemical reagents and custom DNA primers were purchased from Sigma-Aldrich. MS solvents were all high-performance liquid chromatography (HPLC) grade and purchased from Thermo Fisher Scientific. Eppendorf tubes used for MS were Ultra High recovery Eppendorf tubes (STARLAB).

### Cell Culture, transient transfection and incubation

All experiments were performed using three independent experiments (biological replicates). One million GFP SK-N-AS cells (ECACC catalog no. 94092302) were seeded in 10 cm plates in Dulbecco’s modified Eagle’s media (DMEM) supplemented with 10% (v/v) fetal calf serum, 1% (v/v) nonessential amino acids, and 1% (v/v) penicillin-streptomycin and incubated at 37°C, 5% CO_2_, and 21% O_2_for 24 hours. Cells were tested monthly for mycoplasma infection. Transient transfection was performed 72 hours before experimental use, using polyethylenimine (PEI) 40K MAX, Linear (Polysciences #24765-1). All plasmid stocks were diluted using reduced-serum medium (Opti-MEM, Thermo Fisher) was used to dilute DNA to a final concentration of 10 ng/µL. A stock 1% PEI (w/v) in phosphate buffered saline (PBS) (pH 7.5, adjusted with NaOH) solution was used at a ratio of 4µL PEI: 1 µg total DNA, briefly vortexed and incubated at room temperature for 30 min. The volume of transfection mix added to cultured cells was equivalent to 5% of the total cell culture volume and incubated for 72 h (incubation length shown to induce metastatic phenotype in SK-N-AS cells (16) at 37 °C, 5% CO_2_and either in 21% O_2_or 1% O_2_(Don Whitley H35 Hypoxystation) atmosphere before lysis.

### Chick embryo – CAM assay

CAM implantation was carried out as previously described (16). Briefly, NB cells were harvested and resuspended in serum-free media at a concentration of 1×10^6^ cells/μl. CAM implantation was achieved by transferring 2 μl of the cell suspension into the membrane fold created by careful laceration of a chick embryo at E7. After CAM implantation, eggs were incubated until E14 and scanned for the presence of tumors using a standard stereo fluorescence microscope (Leica M165-FC). Tumors were harvested from the CAM and used for microarray gene expression profiling.

### Microarray Gene Expression Profiling and data analysis

RNA was isolated as described above, and then reverse transcribed, with the cDNA labelled with fluorescent Cy3 dye using the Agilent Low-input Quick Amp Kit (Agilent, UK). cRNA was purified using RNeasy columns (Qiagen, UK) and hybridized overnight to Agilent Human 8×60k Whole Genome microarrays, according to the manufacturer’s protocol. Microarrays were scanned using an Agilent SureScan microarray scanner and processed using Agilent Feature Extraction software. Analysis was performed using R. Probes that did not have a median signal significantly above background (value > 64) were removed. All data were quantile normalized. Probes that could potentially cross-hybridize with chicken cDNA were removed as previously described (19). Batch effects relating to the date the RNA was extracted were removed using the ComBatmethod (20), available in the R package *sva*. Differentially expressed genes were identified by analysis of variance (tumours) comparing hypoxic versus normoxic samples. P-values were corrected for multiple testing error using the False Discovery Rate method of Benjamini and Hochberg (21). Differentially expressed genes had a false discovery rate of <10%.

### Survival Analysis

TARGET (11) and SEQC (22) studies were used selecting non-*MYCN* amplified tumours only. Normalized gene expression data and subject characteristics data from the TARGET study was downloaded from [ocg.cancer.gov]. This expression data “is normalised and summarised using affy tools (rna-sketch) and batch corrected for institution using GLM adjusted for tumour stage and MYCN status”. Normalised gene expression data and subject characteristics data from the SEQC study was downloaded from Gene Expression Omnibus using the accession GSE62564. An additional filter was applied to remove a substantial subset of genes with very low expression: genes with median expression in the lower quartile across all arrays were removed. The smaller TARGET cohort was filtered to include only grade 4 (INSS) tumours due to the low number of lower-grade samples (n = 31). INSS grade was included as a covariate in the analysis of the SEQC cohort, in addition to testing each gene in each grade independently. Survival analysis was performed using the Cox proportional hazard model as implemented by the Cox ph function in the R software package (https://www.r-project.org/). Patients were separated into higher and lower expressing groups by finding the partition, which minimizes the p-value from the log-rank test. Each gene was assessed according to a set of criteria designed to identify those genes that are controlled by hypoxic preconditioning as well as being predictive of outcome. These were: Cox proportional hazards p-value < 0.05 consistency between the high-risk and low-risk groups in terms of which group had the highest expression, absolute log2 fold change (*in vivo* hypoxic preconditioning versus normoxic) > 0.5.

### Cloning of HA-mCherry/mCherry-HA plasmid constructs

All cloning was performed using the In-Fusion Cloning technology (Clontech): HA-mCherry-zDHHC23 (Addgene no.246459), zDHHC23-mCherry-HA (Addgene no. 246460), following manufacturer’s recommended protocols. Plasmid amplification was done using the PureLink HiPure Plasmid Maxiprep Kit (Invitrogen). All cloned plasmids were validated by full construct sequencing (Plasmidsaurus).

### IP and sample preparation for proteomics

Post treatment by transfection and normoxic/hypoxic incubation for 72 hours, cells were lysed in 600 µL lysis buffer (50 mM Tris pH 8.0, 120 mM NaCl, 5 mM EDTA, 0.5% (v/v) NP-40, 1X EDTA-free cOmplete protease inhibitor (Roche) and 1X phosSTOP (Roche). Cells were scraped and supernatant collected into Ultra-High recovery Eppendorf tubes (STARLAB). Lysates were rotated end-over-end for 30 min at 4 °C, before centrifugation at 10,000 *g* for 10 min at 4 °C with the cleared supernatant collected. Pre-complex HA-beads (Thermo Fisher # 13464229) were washed three times in lysis buffer before being added to lysate (5 µL/ 200 µg) and left to rotate end-over-end for 18 hr at 4 °C. Bound complexes were washed three times in lysis buffer, proteins eluted in 2% SDS in 100 mM Tris pH 8.0 volume equal to bead slurry used. Using a MagRack (Cytiva #GE28-9489-64) beads were discarded and supernatant collected. Protein concentration was determined using a BCA protein assay (Thermo Scientific #J63283.QA). For western blotting proteins were then heated 95 °C for 10 min in 4 X Laemmli buffer (250 mM Tris-HCl pH 6.8, 30% (v/v) glycerol, 10% (w/v) SDS, 500 mM DTT, 0.05% (w/v) bromophenol blue). For LC-MS/MS analysis proteins were subject to reduction and alkylation as previously described (23) . The eluent was digested in 10:1 (w/w) trypsin gold (Promega) using the manufacturer’s recommended temperatures for 18 hours with 600 rpm shaking. Samples were split 95:5% and dried to completion under cooled vacuum centrifugation. 95% of the samples were subjected to titanium dioxide (TiO_2_) phosphopeptide enrichment, as described in (23). All dried peptides were solubilized in 20 µl of 3% (v/v) acetonitrile and 0.1% (v/v) TFA in water, sonicated for 10 min, and centrifuged at 13,000g for 15 min at 4°C before liquid chromatography–mass spectrometry analysis (5%: high-low method, 95%: high-high method).

### Liquid chromatography–mass spectrometry (LC-MS) analysis

Peptides were separated by reversed-phase capillary HPLC on an UltiMate 3000 nano system (Dionex) over a 60 min gradient as described in (23). All data acquisition was performed using a Fusion Lumos Tribrid mass spectrometer with higher-energy C-trap dissociation (HCD) fragmentation set at 32% normalized collision energy for 2+ to 5+ charge states. For 5% sample, a high-low method was used; MS1 spectra were acquired in the Orbitrap (120K resolution at a m/z 200) over a m/z range of 350-2000, AGC target = 50%, maximum injection time = 50 ms. MS2 data were acquired in a data-dependent acquisition (DDA) mode, using a ‘top speed’ method with a cycle time of 3 sec. Ion trap MS2 settings were as follows: rapid mode (15K resolution at m/z 200), maximum injection time = dynamic, fragmentation intensity threshold = 1E4. A dynamic exclusion window of 60 s was applied at a 10 ppm mass tolerance. For phosphopeptide enriched samples, a high-high method was used; MS1 spectra were acquired in the Orbitrap (120K resolution at a m/z 200) over a m/z range of 350-2000, AGC target = 50%, maximum injection time = 50 ms. MS2 data was acquired in a DDA mode, using a top speed method with a cycle time of 3 s. MS2 settings were: 30K resolution at m/z 200, maximum injection time = dynamic, fragmentation intensity threshold = 2.5E4. A dynamic exclusion window of 20 s was applied at a 10 ppm mass tolerance.

### MS data analysis

Digested 5% samples were analysed using Proteome Discoverer (PD) v2.4 and searched against the UniProt Human reviewed database (updated weekly). Trypsin enzyme parameters were (cleavage pattern: maximum number of permitted miscleaves): Trypsin (K/R –not P: 2). PD settings are the following: instrument type = ESI-FTICR with fixed modifications = cysteine carbamidomethylation and variable modifications = methionine oxidation, two miscleaves allowed and mass tolerances were MS1 = 10 ppm and MS2 = 0.5 Da. All files were searched separately to create a filtered list that was seen in two out of three replicates, before background subtraction using relevant controls (HA-mCherry/mCherry-HA). A second search combining all files was performed with the minora feature detector node set to calculate peptide abundance (area under the curve) and low abundance resampling imputation of missing values (normalized distribution below the lowest peptide intensity identified) to allow label free quantification (LFQ) and statistical analysis. The background subtracted, seen two out of three replicate list was then used to filter relevant protein identifications in the LFQ and imputed dataset. Data was exported and log2 ratios for 1% O_2_protein intensities/21% O_2_protein intensities were calculated. Fold change (1% O_2_/21% O_2_) and q values were transformed (log2) and imported into a custom R script that colours points red if they have a q value of 5 with their gene name.

Phosphopeptide-enriched samples were analysed using either PD or PEAKS to identify sites of phosphorylation or other PTMs respectively. PEAKS Studio was used as above [using the UniProt Human reviewed database], except for the following adjustments: instrument = Orbi-Orbi, additional variable modifications of phospho S/T/Y and MS2 mass tolerance = 0.01 Da. Proteome Discoverer (PD) in conjunction with MASCOT v2.7 was also used to analyse phosphopeptide enriched data according to (23, 24). A custom database was used for PD analysis of phosphopeptides, as described in (24), which was generated from all identifications from the respective trypsin digested (5%) sample that was analysed by PD on the UniProt Human reviewed database (updated weekly) with fixed modification = carbamidomethylation (C), variable modification = oxidation (M), instrument type = electrospray ionization–Fourier-transform ion cyclotron resonance (ESI-FTICR), MS1 mass tolerance = 10 ppm, and MS2 mass tolerance = 0.5 Da. Phosphopeptide analysis PD settings are the following: instrument type = ESI-FTICR, variable modifications = carbamidomethylation (C), oxidation (M) and phospho (S/T/Y/D/E/K/R/H/C). MS1 mass tolerance = 10 ppm, MS2 mass tolerance = 0.01 Da and the ptmRS node on, filtered to a ptmRS score > 98.0. All data from each analysis pipeline were filtered to a 1% FDR at the PSM (peptide spectrum match) level.

### Bioinformatics

#### Gene Ontology - functional enrichment analysis

DAVID Bioinformatics Resources v2024q4 (25) was used to understand differences in our transcriptomic data and zDHHC23 binding partners/palmitoylated targets as a function of O_2_tension (q value < 0.05), using a background database of *Homo sapiens*. Data was filtered so that GOterms: Molecular Function (MF), Cellular Compartment (CC) and Biological Process (BP), and KEGG Pathway annotations were maintained. P-values were adjusted using the Benjamini-Hochberg method. Data visualization was performed using a custom R script and labelling amended in InkScape v1.2.

#### Comparison with BioGRID data

BioGRID v4.4 (26) was accessed on May 27, 2025. Interactors of zDHHC23 identified exclusively through affinity capture–mass spectrometry (70 proteins; Supplementary File 6) were compared to the interactors identified in our study. Data visualization was performed using a custom R script, highlighting known BioGRID interactors as red dots on the LFQ volcano plot.

#### Phylogenetic analysis

Human zDHHC23 (Q8IYP9) protein sequence was used as a starting point to create phylogenetic trees following guidance from (27). Briefly, the selected protein sequences were searched using BLAST (Basic Local Alignment Search Tool) to identify the top 500 homologous sequences. All “partial” and “unknown protein”–labelled proteins were removed. Genus-species names were converted to “common names” using the Taxize plugin for R (28) [using the Global Names Resolver webpage (https://resolver. globalnames.org/)]. Sequences were aligned using the multiple sequence alignment tool MUSCLE (29), and a phylogenetic tree was produced using MEGA7 (30) with 500 bootstrap replicates. Alignments were reordered to match the phylogenetic tree order and viewed using ClustalX (31).

#### STRING protein-protein interaction analysis

STRING v12, accessed at https://string-db.org was used for network analysis, with proteins entered as a list using their UniProt accessions and searched against the *Homo sapiens* database. In all cases, data were filtered so that only interactions identified in experimental, or database-level evidence were included at the highest confidence, and any nodes with less than three connections were removed for viewing. KMEANS clustering was used to identify clusters of proteins with shared functions, the number of clusters set was determined empirically for each network. Networks generated in STRING were exported as a tab-separated values (.tsv) file and imported into Cytoscape v3.9.1 (32) for editing and PowerPoint to customize the overall network presentation.

#### Florescent confocal microscopy for sub-cellular localization

SK-N-AS cells were seeded in 12 well glass bottom dishes (Cellvis #P12-1.5H-N) and transfected as described above, followed by 72 hr incubation in either 21% or 1% O_2_. Media was removed and cells washed with PBS. Cells were then fixed with 1 ml 4% (w/v) paraformaldehyde at room temperature for 10 min before a wash in PBS followed by a Thermo Fisher Hoechst nuclear stain (#62249, 1 in 1000) added for 3 min before a final wash with PBS. Confocal images were acquired using a LSM780 Zeiss Microscope with Zen 2012 software, equipped with 488 nm, 561 nm and 405 nm filters at a magnification of 63X.

## Results

### Transcriptomic profiling of hypoxia preconditioned metastatic neuroblastoma tumours

To interrogate the molecular basis of neuroblastoma (NB) metastasis independent of *MYCN* amplification, we utilised our previously established chick embryo xenograft model, in which hypoxic preconditioning (1% O_2_) reliably induces metastatic dissemination of NB cells across multiple organ systems (16). This phenotype was consistently observed in both *MYCN*-amplified (SK-N-BE(2)C) and non-*MYCN* amplified (SK-N-AS, 11q-deleted) cell lines, confirming that hypoxia-driven metastatic competence is not restricted to *MYCN* status. Building on these investigations, we preconditioned SK-N-AS cells under normoxic or hypoxic conditions prior to implantation onto the chorioallantoic membrane (CAM) of chick embryos. Following tumour formation, primary xenografts were harvested and subjected to microarray-based gene expression profiling to dissect the transcriptional landscape underpinning this phenotype (Figure 1A, Supplementary File 1). Comparative analysis revealed a robust hypoxia-associated gene signature comprising 398 differentially expressed genes (DEGs), with 322 transcripts significantly upregulated and 76 downregulated in hypoxia-preconditioned (metastatic) tumours relative to normoxic (non-metastatic) controls (Figure 1B).

**Fig 1.**
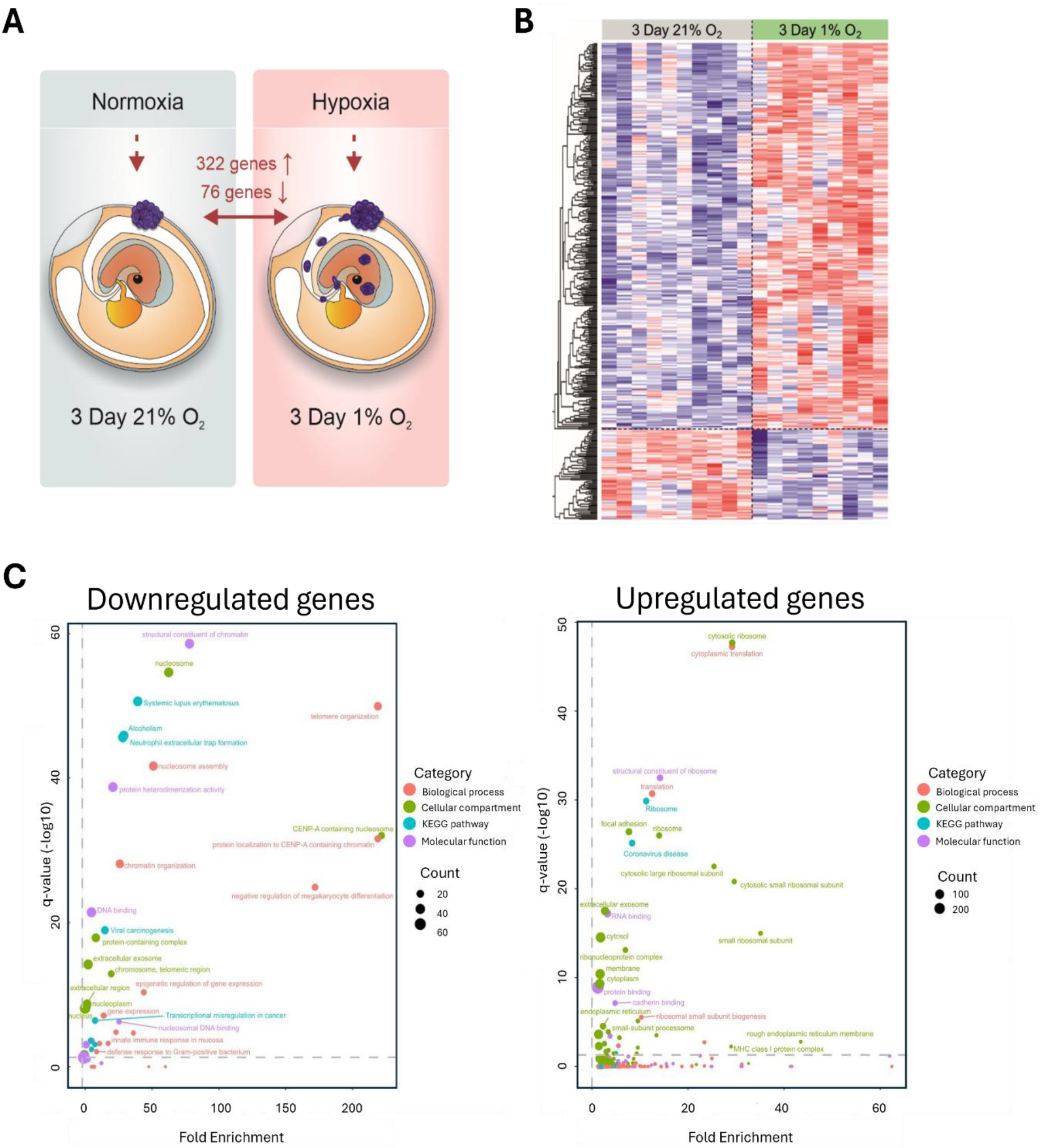
Transcriptomic profiling of tumors after hypoxic preconditioning. **A** Schematic of the experimental setup to define hypoxia-dependent differential gene expression profiles. SK-N-AS cells were grown in vivo in normoxia (3 days in 21% O_2_) or hypoxia (3 days in 1% O_2_) and extracted RNA subjected to microarray gene expression profiling. The number of significantly (Benjamini-Hochberg (BH) adjusted p-value <0.1) differentially expressed genes between conditions is indicated. **B** Heatmap representation with hierarchical clustering of the differentially expressed genes. Bar plots adjacent to the heatmap show over-represented Gene Ontology and KEGG terms in the differentially expressed genes. Bar values indicate the number of genes matching a given term. Heatmap expression levels have been standardised in a row-wise manner (z-score) to visualise genes of different expression levels on the same plot. **C** GO enrichment analysis was performed in DAVID using the list of either significantly downregulated (left) or upregulated (right) genes. P-values were adjusted for multiple hypothesis testing using BH correction and plotted against fold enrichment. The horizontal line represents a BH adjusted p-value of 0.05. Points are sized according to the number of proteins within each enriched term and coloured according to the enrichment category.

Functional characterisation of the down-regulated genes were represented by terms such as *telomere organisation* (q-value = 1.1E-50), *nucleosome assembly* (q-value = 2.2E-42), *neutrophil extracellular trap formation* (q-value = 2.4E-46) and *transcriptional misregulation of cancer* (q-value = 3.9E-07) (Figure 1C, Supplementary File 2). Up-regulated genes were represented by terms such as *translation* (q-value = 2.0E-31), *RNA binding* (q-value = 6.1E-18), and *cadherin binding* (q-value = 7.3E-08).

Encouragingly, this hypoxic gene signature included classic hypoxia target genes such as *AK4* (1.8-fold, q-value = 6.8E-03), *ADM* (1.7-fold, q-value = 2.8E-02), *NDRG1* (1.7-fold, q-value = 5.1E-02), *SLC25A29* (1.3-fold, q-value = 8.3E-02), *PDGFRB* (1.2-fold, q-value = 8.0E-02), *DHX35* (0.8-fold, q-value = 7.6E-02) and *CDKN1B* (0.7-fold, q-value = 5.3E-02), alongside genes involved in detachment, invasion and intravasation such as *ADAMTS9* (1.4-fold, q-value = 2.3E-02), *TIMP3* (1.3-fold, q-value = 6.4E-02) and *SNAI* (0.8-fold, q-value = 7.0E-02), and cell adhesion such as *ITGA7* (1.6-fold, q-value = 6.3E-02). Interestingly, a gene that regulates the stability of MYCN in neuroblastoma (*MYCNOS*) was downregulated in hypoxia (0.7-fold, q-value = 2.8E-02), while a number of novel genes were substantially elevated, including *UBC* (1.9-fold, q-value = 2.1E-04), *MROH6* (2.0-fold, q-value = 7.7E-02) and *zDHHC23* (1.9-fold, q-value = 8.8E-02).

Promisingly, 20 of our DEGS were also identified in another study which focused on differentially expressed proteins from metastatic tissue (ovarian and liver) derived from a xenograft mouse model of human NB (33). Interestingly, this included nucleoside diphosphate kinase B (NME2), a known histidine kinase (34) reported to act as both a suppressor of metastasis and of apoptosis in different cancer environments (35, 36). In agreement with the data from Hanel *et al* (33) which reported elevated NME2 levels in both ovarian and liver metastasis, we also identified upregulation of *NME2* (1.4-fold; q-value = 4.8E-02) in our NB cells upon hypoxic preconditioning, suggesting this may be worthy of further exploration for use as a NB metastatic marker.

### zDHHC23 is predictive of survival in non-MYCN amplified patients

To evaluate whether the hypoxia-regulated gene signature identified using our *in vivo* model reflects clinically aggressive NB, we performed a survival analysis of the 398 DEGs using two independent NB patient cohorts: TARGET (11) and *SEQC* (22) (see Methods). Of these, 59 transcripts were significantly associated with event-free survival (EFS) time (p < 0.05), suggesting prognostic relevance in non-*MYCN* amplified disease. To prioritise candidates most likely to represent robust markers of hypoxia-driven metastatic potential, we applied stringent filtering criteria based on expression concordance, fold change, and survival association across both datasets. Seven genes *(zDHHC23, MALAT1, AP2S1, UCP3, RPS21, RPS2*, and *SLC7A5*) met all criteria and were consistently predictive of poor outcome. An additional three genes (*MROH6, IL6ST, CCDC177*) demonstrated significant survival association in one cohort and exhibited high differential expression (log_2_FC > 1) in the other (Table 1).

**Table 1.** Survival Analysis. Genes were selected according to a set of criteria designed to identify those that were likely to be controlled by hypoxic preconditioning as well as being predictive of patient outcome in independent datasets. Choice of genes is based on fold change (FC), hazard ratio (HR) and p-value from SEQC and TARGET databases. P-values that do not pass the 0.05 cut-off are in italics.

| Gene | Name | FC | TARGET |  | SEQC |  |
| --- | --- | --- | --- | --- | --- | --- |
|  |  |  | HR | p-value | HR | p-value |
| MROH6 | Maestro Heat Like Repeat Family Member 6 | 2.01 | 2.00 | 1.85 E-03 | 1.58 | 5.38 E-02 |
| ZDHHC23 | Zinc Finger DHCC-Type Containing 23 | 1.88 | 2.96 | 1.00 E-06 | 2.20 | 2.67 E-04 |
| RPS2 | Ribosomal Protein S2 | 1.68 | 1.58 | 3.44 E-02 | 1.56 | 2.95 E-02 |
| SLC7A5 | Solute Carrier Family 7 Members S | 1.64 | 1.55 | 4.17 E-02 | 1.84 | 2.49 E-02 |
| RPS21 | Ribosomal Protein S21 | 1.55 | 1.65 | 2.02 E-02 | 1.78 | 8.45 E-03 |
| AP2S1 | Adaptor Related Protein Complex 2 Subunit Sigma 1 | 1.42 | 1.81 | 1.28 E-02 | 1.94 | 8.59 E-04 |
| MALAT1 | Metastasis Associated Lung Adenocarcinoma Transcript 1 | 0.65 | 1.81 | 6.55 E-03 | 1.77 | 1.80 E-02 |
| UCP3 | Uncoupling Protein 3 | 0.56 | 1.78 | 2.11 E-02 | 2.09 | 1.41 E-03 |
| IL6ST | Interleukin 6 Signal Transducer | 0.49 | 1.50 | 8.05 E-02 | 2.03 | 7.92 E-03 |
| CCDC177 | Coiled-Coil Domain Containing 177 | 0.28 | 1.34 | 1.35 E-01 | 2.09 | 8.17 E-03 |

To contextualise these findings, we compared our hypoxic gene signature with a previously published murine model of non-*MYCN* amplified NB metastasis based on intracardiac injection of SK-N-AS cells (37). Six genes overlapped between datasets including *ADM* (1.7-fold, q-value = 2.8E-02), *COL1A1* (1.8-fold, q-value = 8.8E-03), *ZNF865* (1.6-fold, q-value = 2.7E-03), *ZNF814* (1.3-fold, q-value = 9.2E-02), *ZNF394* (0.8-fold, q-value = 9.2E-02), and *CDKN1B* (0.7-fold, q-value = 5.3E-02). However, none met our prognostic thresholds due to low hazard ratios and modest fold changes.

Among the top candidates, *zDHHC23* emerged as a compelling prognostic marker. Elevated *zDHHC23* expression was associated with significantly reduced survival probability in both low-grade (I–III) and high-grade (IV) patients (Figure 2, Supplementary Figure 1). Notably, low-grade patients with high *zDHHC23* expression exhibited survival outcomes comparable with high-grade cases, underscoring its clinical relevance. This stratification was independent of *MYCN* amplification status and suggests that zDHHC23 may serve as a more universally applicable biomarker of NB aggressiveness. These findings are consistent with emerging evidence implicating zDHHC23 in other malignancies, including glioblastoma multiforme (38) and B-cell neoplasms (39), where it has been linked to stem cell maintenance and malignant progression. Most recently, zDHHC23 has been proposed as a prognostic marker in osteosarcoma (40), further supporting its role in regulating cellular programs exploited during tumour evolution.

Of the remaining 8 hypoxia-regulated genes predictive of outcome, *ASP2S1, UCP3, IL6-ST* and *CCDC177* demonstrated good stratification of both low- and high-grade patients albeit to a lesser extent than *zDHHC23* (Table 1). *MALAT1*, like *MROH6*, was predictive only for low-grade tumours (Supp. Figure 1). Based on the strength of its prognostic association and hypoxia responsiveness, we next sought to characterise the molecular functions of zDHHC23 in non-*MYCN* amplified SK-N-AS cells, with the aim of better defining its potential roles in metastatic signalling.

### Hypoxia regulates the interaction network of zDHHC23

To elucidate the protein interaction network of zDHHC23, a relatively understudied palmitoyltransferase, and assess its modulation by oxygen tension (a key determinant of metastatic behaviour), we performed mass spectrometry (MS)-based label free quantification (LFQ) following HA-based immunoprecipitation of transiently expressed zDHHC23 in SK-N-AS cells under normoxic (21% O_2_) or hypoxic (1% O_2_) conditions. Constructs bearing either HA-mCherry- or -mCherry-HA tags were used to control for potential tag-orientation effects. Only proteins identified in ≥2 of 3 replicates per construct were retained for downstream analysis to ensure robustness.

Across conditions, 446 high-confidence zDHHC23 interactors were identified, of which 81 were significantly more abundant in normoxia (Figure 3A, Supplementary File 5). Integration with BioGRID interactome data revealed two common protein binding partners, POF1B and UPF1, with only POF1B being statistically significantly enriched in our normoxic dataset (Figure 3A, red dots; Supplementary File 6).

**Figure 3.**
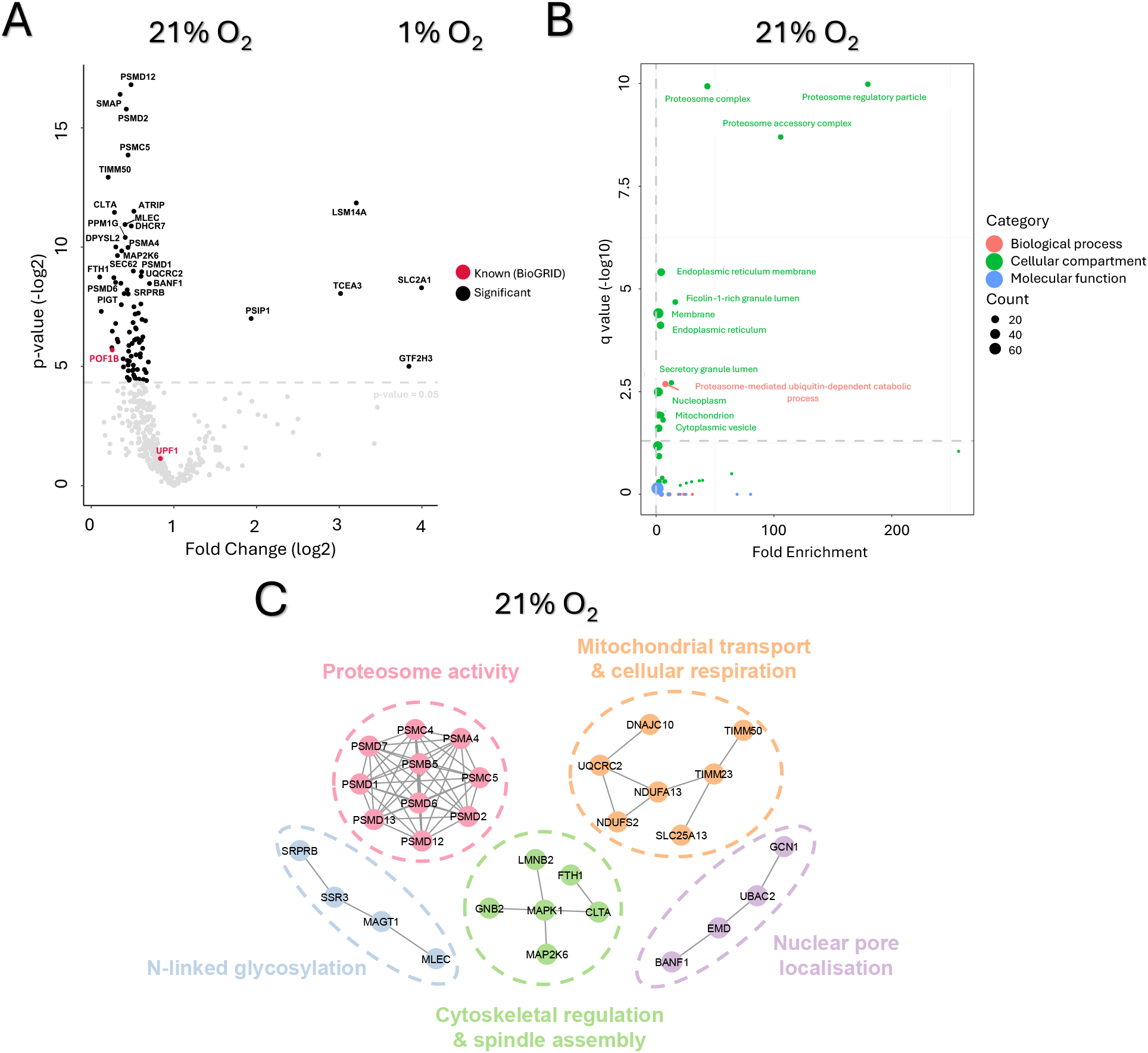
Interactome of zDHHC23 is notably decreased in hypoxia. **A)** Volcano plot depicting statistically significant changes in the relative abundance (log2 fold change) of zDHHC23 interacting proteins in normoxia (21% O_2_) compared with hypoxia (1% O_2_). The horizontal line represents an adjusted p-value of 0.05 with known interactors identified via BioGRID (24) colored red and significant proteins colored black. Point size is scaled to the total number of peptide spectrum matches per protein. **B)** GO enrichment analysis of the significant oxygen tension-dependent interactors (identified in at least 2 replicates) was performed in DAVID. P-values were adjusted for multiple hypothesis testing using the Benjamini-Hochberg correction and plotted against fold enrichment. The horizontal line represents an adjusted p-value of 0.05. Points are sized according to the number of proteins within each enriched term and colored according to the enrichment category. **C)** Interaction network of proteins significantly elevated in normoxia, with nodes organized and colored nodes into functional groups/complexes. The original network was generated in STRING to include interactors with experimental or database evidence of medium confidence (interaction score 0.400).

A prominent feature of the zDHHC23 hypoxic interactome was the reduction of proteasome-associated proteins. Of the 28 subunits of the 20S core particle, two subunits (PSMA4 and PSMB5) were significantly reduced, while three others (PSMA2, PSMB3, and PSMB4) showed non-significant decreases. Similarly, association of zDHHC23 with 10 subunits of the 19S regulatory particle was reduced in hypoxia, with eight (PSMC4, PSMC5, PSMD1, PSMD2, PSMD6, PSMD7, PSMD12, and PSMD13) reaching statistical significance. PSMD12 emerged as the most significantly affected interactor (p-value = 8.7E-06), reducing over 2-fold in hypoxia. Gene Ontology analysis highlighted proteasome-related terms as dominant features (Figure 3B), while STRING network analysis revealed a tightly interconnected cluster of regulated proteasomal proteins (Figure 3C).

Hypoxia also disrupted interactions with TIMM50 and TIMM23, key components of the TIM23 complex responsible for translocating nuclear-encoded proteins across the inner mitochondrial membrane (41, 42). Elevated expression of TIMM50 in multiple tumor types has been reported previously, where it is thought to promote tumor progression by enhancing mitochondrial function and facilitating metabolic adaptation (43, 44). STRING analysis identified a cluster of seven dysregulated proteins involved in mitochondrial transport and respiration (Figure 3C), implicating zDHHC23 in oxygen-sensitive regulation of mitochondrial function. TIMM50, previously linked to tumour progression via enhanced mitochondrial activity, was notably downregulated under hypoxia.

In addition, interaction with four nuclear pore-associated proteins (BANF1, EMD, UBAC2, GCN1) was disrupted in hypoxia (Figure 3C). Each has been implicated in cancer-related processes: BANF1 correlates with poor prognosis in gastric cancer (45), UBAC2 is overexpressed in bladder cancer (46), GCN1 is elevated across multiple tumour types (47-49), and EMD has been linked to breast cancer metastasis (50). STRING analysis confirmed their functional clustering (Figure 3C), suggesting that zDHHC23 may also interface with nuclear transport machinery in an oxygen-dependent manner.

Only five proteins were significantly elevated in the hypoxic interactome (Figure 3A, Supplementary File 5). Although these proteins do not share a single overarching function, each is likely to play a unique role relevant to metastatic NB biology. LSM14A, the most significantly upregulated interactor (p-value = 2.7E-04; 3.2-fold), is essential for P-body formation and mitotic spindle assembly, and has been implicated in chromosomal instability and tumour aggressiveness (51) (52). SLC2A1, a HIF-1 target facilitating glucose uptake for anaerobic glycolysis, showed the highest fold change (∼4-fold; p-value = 3.2E-03), consistent with metabolic reprogramming under hypoxia (53). Additional hypoxia-enriched interactors included PSIP1, TCEA3, and GTF2H3, proteins involved in transcriptional elongation, cell cycle regulation, and genomic stability. PSIP1, a chromatin-associated protein, promotes breast cancer progression and drug resistance (54-56), while the transcriptional elongation factor TCEA3 has been shown to induce apoptosis in multiple tumour types (57-59). GTF2H3, a core component of the TFIIH complex, is essential for transcription initiation and nucleotide excision repair (60), implicating it in tumour evolution and genomic maintenance (61).

### PTM status of zDHHC23 is minimally regulated by hypoxia

PTMs are critical modulators of protein function, influencing enzymatic activity, subcellular localisation, and protein–protein interactions. Under differential oxygen tension, PTMs such as phosphorylation, hydroxylation, and oxidation are frequently altered, contributing to cellular adaptation and metastatic progression (62, 63). Members of the zDHHC family of palmitoyltransferases are known to undergo diverse PTMs, including phosphorylation, acetylation, ubiquitination, methylation and S-acylation (64), yet the oxygen-dependent regulation of these modifications remains poorly defined.

To investigate whether hypoxia modulates the PTM landscape of zDHHC23, we performed phosphopeptide enrichment on tryptic digests of immunoprecipitated zDHHC23 from SK-N-AS cells cultured under normoxic (21% O_2_) or hypoxic (1% O_2_) conditions. LC-MS/MS analysis identified four sites of modification, including two known phosphorylation events: condition-independent phosphorylation of Ser206 (pSer206) and normoxia-specific Ser252 phosphorylation (pSer252) (65) (Figure 4A, Supplementary Figure 2 and File 3). Additionally, we detected hypoxia-specific oxidation of Asp44 and oxygen-independent oxidation of Trp260.

**Figure 4.**
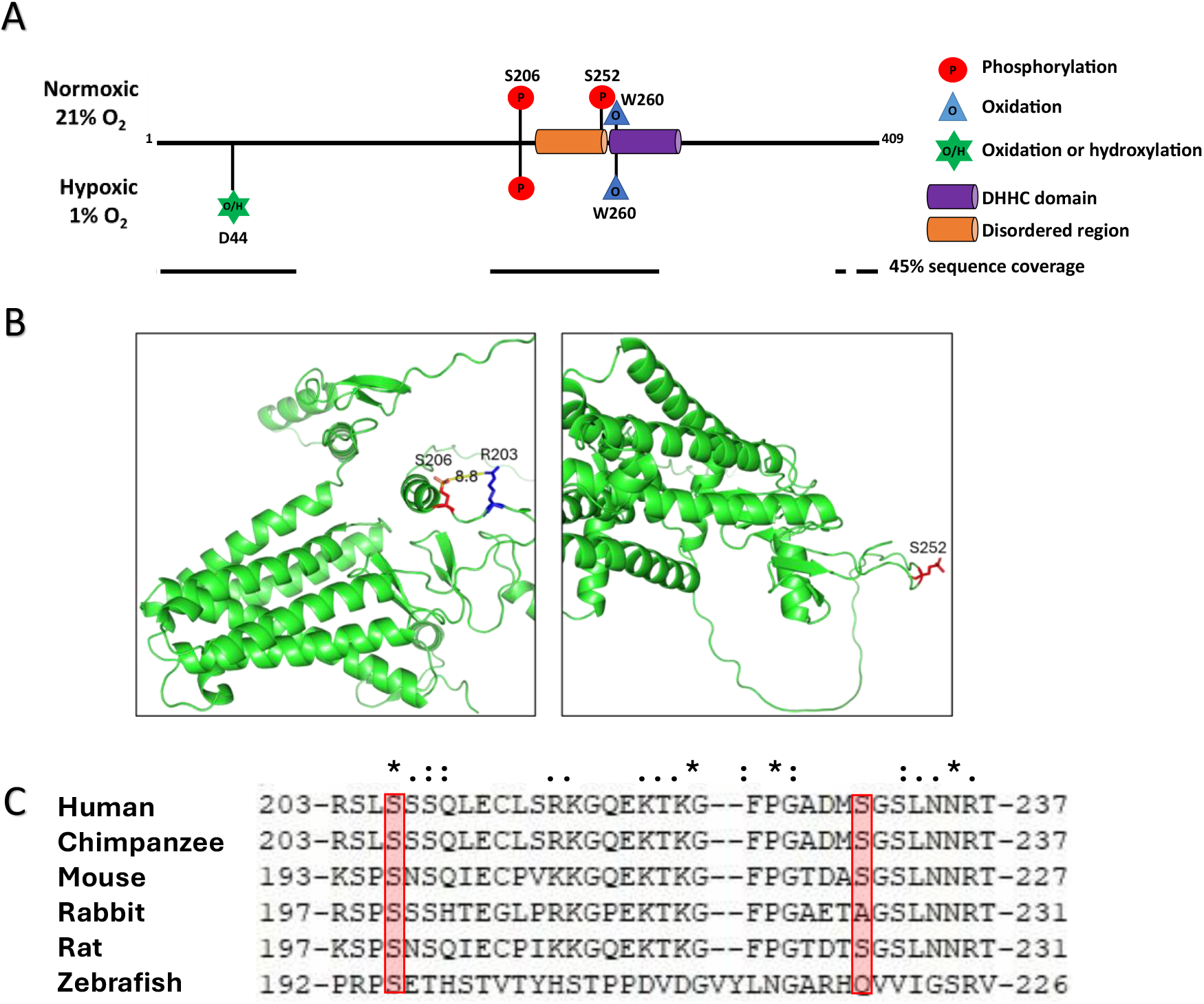
PTM sites identified on zDHHC23 expressed in SK-N-AS cells cultured in either 21% O_2_or 1% O_2_. **A)** Schematic sequence map of zDHHC23. Confidently identified PTMs are mapped to the protein and coloured dependent on the type of modification. LC-MS/MS sequence coverage is depicted below the protein schematic. Data presented are from three independent biological replicates for each condition, with all sites seen in at least 2 of 3 replicates. PTMs with a black outline represent modifications observed at least four times on PhosphoSitePlus. **B)** AlphaFold 3 structure prediction of zDHHC23 *in silico* phosphorylated at either S206 (left) or S252 (right) Green: zDHHC23, Red: phosphorylation site, Blue: possible S206 interacting residue R203 and D44 interaction with R76 that is likely disrupted upon oxidation. Distance was measured using PyMOL distance wizard. **C)** Sequence alignments of zDHHC23 in human, chimpanzee, mouse, rabbit, rat and zebrafish (common *in vivo* research models) in MUSCLE multiple sequence alignment and viewed in Clustal X; (*) - identical residue; (:) - similar residue properties; (.) - less similar residue properties; (space) - no similarity in residues, properties as determined by Clustal X. Red box highlights the two phosphorylation sites.

By mapping the sites of phosphorylation onto the predicted AlphaFold 3 structure of zDHHC23 using PyMOL (PyTMs plugin) (66, 67), we were able to explore the structural implications of these modifications. Interestingly, pS206 localised 8.8 Å distal from Arg203, suggesting the potential formation of a stabilising salt bridge. In contrast, pS252 was situated within a disordered loop adjacent to a predicted *C*-terminal transmembrane anchor, implicating it in membrane association or trafficking (Figure 4B).

Evolutionary conservation analysis (68) revealed that Ser206 is highly conserved across commonly used *in vivo* models, whereas Ser252 exhibited lower conservation, with the corresponding residues being Ala or Glu in rabbit and zebrafish, respectively. Arg203, proximal to pSer206, was also largely conserved across species, further supporting its structural relevance (Figure 4C).

Using the PhosphoSitePlus kinase prediction tool (69), we identified putative regulatory kinases for both phosphorylation sites (Supplementary File 4). This analysis suggested that Ser206 phosphorylation is likely catalysed by a member of the CAMK family, with MARK2 (a known regulator of neuronal cell polarity and migration (70, 71)) being the highest scoring. Ser252 was predicted to be a substrate of CMGC family kinases, with eight of the top ten candidates being cyclin-dependent kinases (CDKs), and CDK3 emerging as the most likely regulator. These findings suggest that pSer252 may be subject to cell cycle-dependent regulation, potentially disrupted under hypoxic conditions (72).

### zDHHC23 subcellular localisation is independent of changes in O_2_tension

Given the zDHHC23 interactome data and the potential oxygen-dependent roles in cytoskeleton regulation, we examined the subcellular localization of exogenously expressed mCherry-tagged zDHHC23 in SK-N-AS cells using confocal fluorescence microscopy (Figure 5).

**Figure 5.**
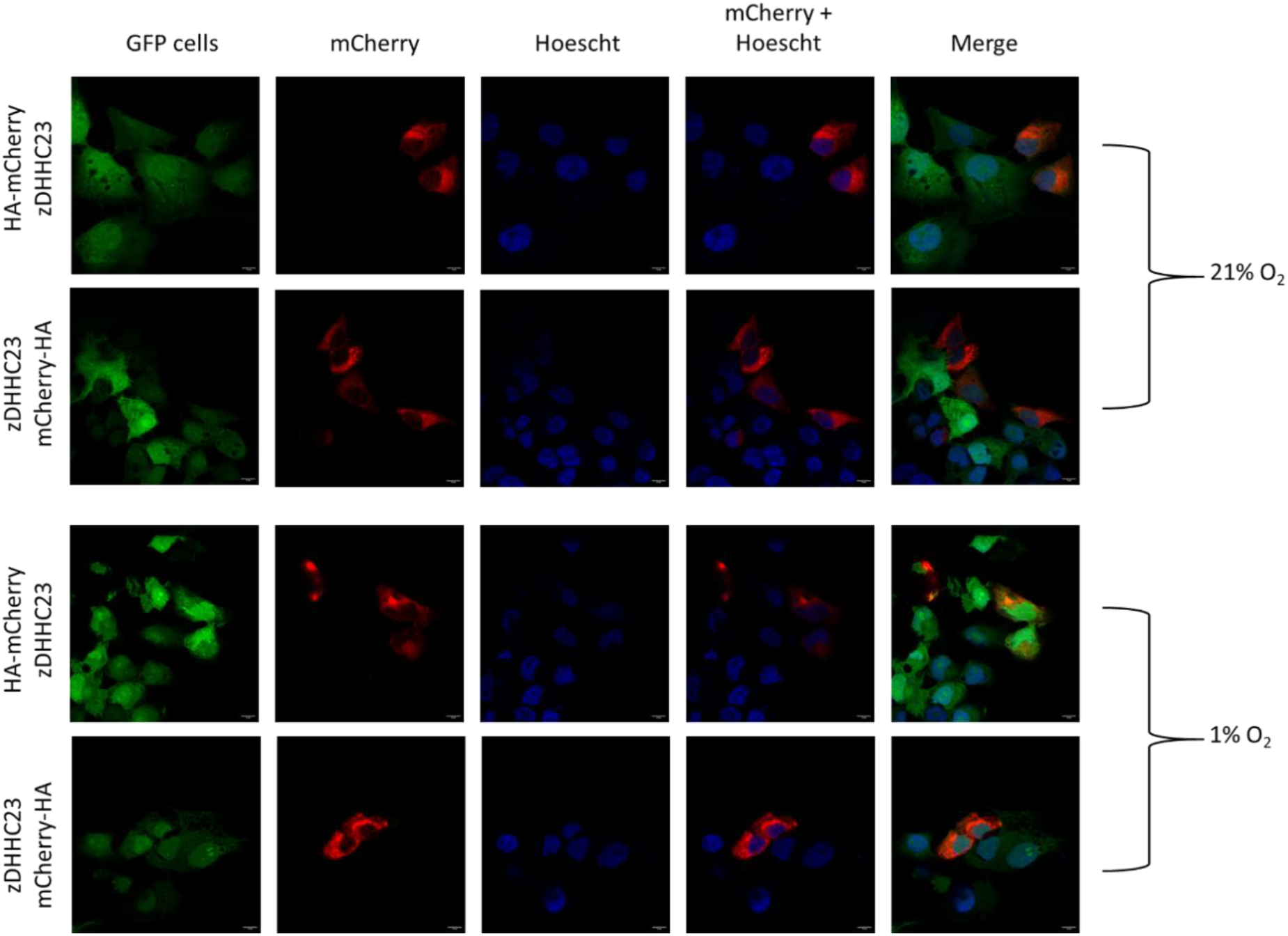
Exogenous zDHHC23 cell localisation in 21% and 1% O_2_. A representative sample of GFP SK-N-AS cells transfected using PEI with N- and C-terminal plasmid constructs and incubated at 21% or 1% O_2_for 72 h post transfection. Cells were fixed with 4% PFA, nuclei stained with Hoechst, and imaged by confocal microscopy. Fluoresence images (405 nm, 488 nm, 587 nm) 63x magnification obtained on Zeiss LSM780. Scale bar = 10 µm.

Across both normoxic (21% O_2_) and hypoxic (1% O_2_) conditions, zDHHC23 consistently exhibited cytoplasmic localisation, with exclusion from the nucleus irrespective of oxygen tension or tag configuration. In addition to diffuse cytoplasmic distribution, peri-nuclear accumulation was also observed, a pattern characteristic of Golgi-associated proteins. This localisation is consistent with previous reports implicating zDHHC family members in Golgi-resident palmitoylation activity and trafficking regulation (73).

## Discussion

Accurate risk stratification at diagnosis is essential for NB patients to receive appropriate treatment. Several transcriptomic studies have hypothesized potential markers for metastatic neuroblastoma group stratification in both *MYCN* (74) and non-*MYCN*-amplified NB (75, 76) which crucially is associated with relapse and death in most patients, but none have yet proven to be efficacious in clinic. Using a chick embryo xenograft model, which we have previously demonstrated can elicit a clear metastatic phenotype following hypoxic preconditioning of tumor cells prior to CAM implantation (16), we have identified hypoxia-induced transcriptional and proteomic changes that underpin metastatic competence in non-*MYCN* amplified neuroblastoma (NB). Transcriptomic profiling revealed a distinct hypoxia-associated gene signature, enriched for pathways involved in translation, RNA binding, and cadherin-mediated adhesion, while downregulated genes were associated with telomere organisation and nucleosome assembly, hallmarks of chromatin remodelling and genomic instability.

Among the differentially expressed genes, *zDHHC23* emerged as a novel compelling candidate biomarker. Elevated expression of *zDHHC23* was significantly associated with poor event-free survival across two independent NB patient cohorts (TARGET and SEQC), and notably stratified outcomes in both low- and high-grade disease. This prognostic relevance was independent of *MYCN* status, suggesting that *zDHHC23* may serve as a universal marker of NB aggressiveness. The clinical significance of zDHHC23 is further supported by its implication in other malignancies, including glioblastoma, B-cell neoplasms, and osteosarcoma, where it has been linked to stem cell maintenance and malignant progression (38-40).

Proteomic analysis of zDHHC23 binding partners revealed an extensive and previously undefined interactome. Of direct relevance to this study, this interaction network was subject to dynamic remodelling under hypoxic conditions. Hypoxia disrupted associations with proteasome subunits, mitochondrial transport proteins (e.g., TIMM50, TIMM23), and nuclear pore components, suggesting that zDHHC23 may act as a central node integrating oxygen-sensitive signalling pathways. The reduction in proteasome-associated proteins under hypoxia aligns with previous reports of proteasomal suppression during metastatic transition, potentially contributing to altered protein turnover and stress adaptation. Indeed, the 20S core particle has previously been linked to NB through co-translational cleavage of nascent proteins that mediate neuronal communication (77).

Conversely, hypoxia enriched interactions with proteins involved in metabolic reprogramming (SLC2A1), chromosomal instability (LSM14A), transcriptional elongation (PSIP1, TCEA3), and DNA repair (GTF2H3). These findings implicate zDHHC23 in diverse cellular processes critical for tumour progression, including glycolytic shift, mitotic fidelity, and genomic maintenance. Notably, the upregulation of SLC2A1, a canonical HIF-1 target, reinforces the metabolic dimension of hypoxia-driven NB aggressiveness.

Despite these extensive changes in the zDHHC23 interactome, the PTM landscape of the protein was minimally affected by oxygen tension. Phosphoproteomic analysis identified two phosphorylation sites (Ser206 and Ser252), with Ser252 being normoxia-specific and potentially regulated by CDK3. Structural modelling suggested functional relevance of these sites, particularly Ser206, which may contribute to protein stability via salt bridge formation. However, the limited PTM modulation under hypoxia suggests that zDHHC23 function is primarily governed by its interaction network rather than direct modification, but incomplete sequence coverage of zDHHC23 in these studies may mean that we are missing a key, yet to be identified, PTM.

Finally, subcellular localisation confirmed that zDHHC23 resides predominantly in the cytoplasm with some peri-nuclear enrichment, consistent with Golgi-associated palmitoylation activity. This localisation pattern remained unchanged under hypoxic conditions, indicating that oxygen-dependent functional shifts are likely mediated primarily through altered complex formation rather than spatial redistribution.

In summary, our findings show that zDHHC23 is associated with hypoxia-responsive changes during metastatic signalling in non-*MYCN* amplified NB. Its prognostic value, dynamic interactome, and involvement in key oncogenic pathways underscore its potential as a biomarker and therapeutic target. Future studies should explore the mechanistic basis of zDHHC23-mediated palmitoylation events and their contribution to NB progression, and assess its utility in clinical stratification and treatment response.

## Data Availability

The mass spectrometry proteomics data have been deposited to the ProteomeXchange Consortium via the PRIDE partner repository with the dataset identifier PXD064446 and 10.6019/PXD064446.

## Acknowledgements

Sally Oswald was funded jointly by the University of Liverpool and Alder Hey Children’s Charity and Oncology Fund (Reference number TF8838). Mass spectrometry platforms used in this study were support by funding from the Biotechnology and Biological Sciences Research Council (BBSRC, BB/R000182/1; BB/M012557/1). The confocal system used in this work was funded by MRC grant number MR/K015931/1. The authors gratefully acknowledge A.H. for her essential contributions to the transcriptomics presented in this manuscript, Francesco Falciani with assistance with data analysis supervision, and Tilly for assistance with bioinformatics analysis. Special thanks to Richard Scheltema and Andris Jankevics for providing access to their spectral annotation software. Microscopy data were acquired at the Centre for Cell Imaging (CCI) at the University of Liverpool.

## Author Contributions

Conceptualization: VS, BP, CE, IP; Methodology: SO, LD, DB, CE, IP; Investigation: SO, LD, KC, PB; Visualization: SO; Funding acquisition: VS, CE, BP, SO; Supervision: CE, IP, BP, VS; Writing – original draft: SO, CE; Writing – review & editing: SO, CE, VS, IP, LD, BP, KC.

## Additional Information

Supplementary Figure 1 – Survival Analysis in the TARGET and SEQC NB patient cohorts

Supplementary Figure 2 - zDHHC23 PTM Annotated Spectra

Supplementary File 1 – Microarray Data

Supplementary File 2 - Gene Ontology Analysis of Upregulated and Downregulated Genes

Supplementary File 3 - zDHHC23 PTM Identification

Supplementary File 4 - zDHHC23 PTM Kinase Analysis

Supplementary File 5 - zDHHC23 Label Free Quantification

Supplementary File 6 – zDHHC23 BioGRID Dataset

Supplementary File 7 - Gene Ontology Analysis of Statistically Significantly Binding Partners in 21% O2

