## Supplementary figures and images for "Hypoxia-induced regulation of zDHHC23 opens avenues for new biomarkers for NON MYCN-amplified neuroblastoma"

### Supplementary Figure 1

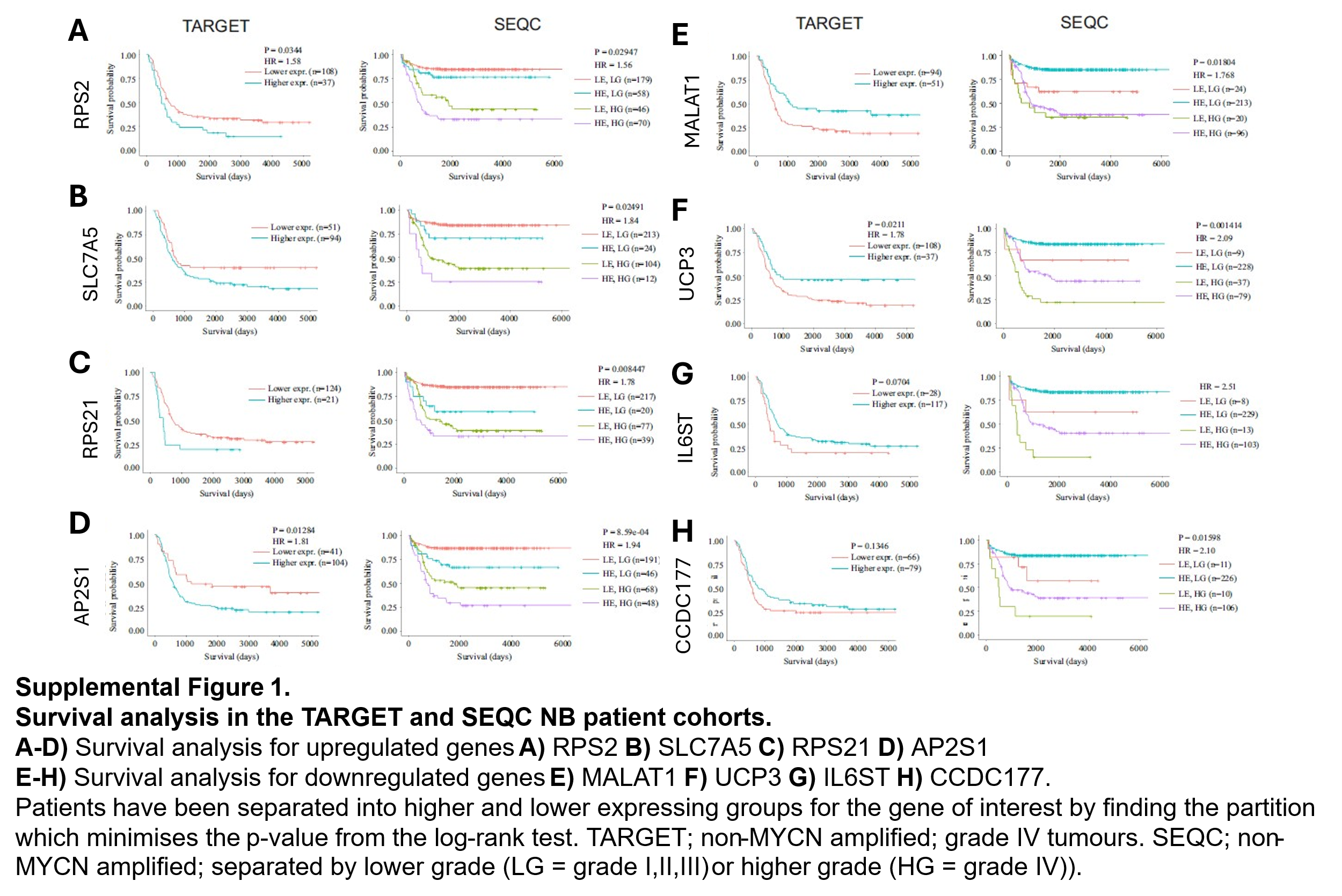
