## Supplementary Figure 2 for "Hypoxia-induced regulation of zDHHC23 opens avenues for new biomarkers for NON MYCN-amplified neuroblastoma"

**A**

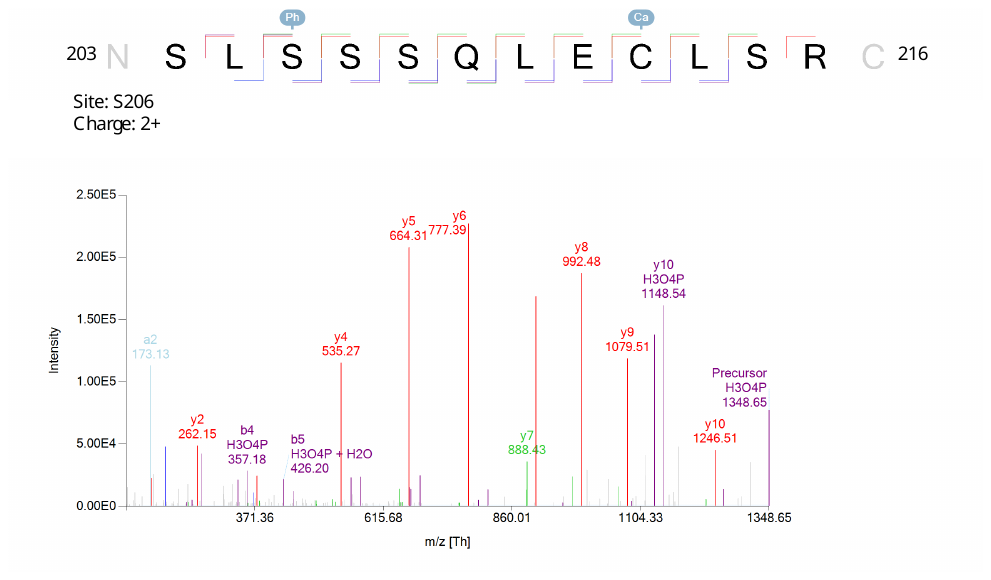

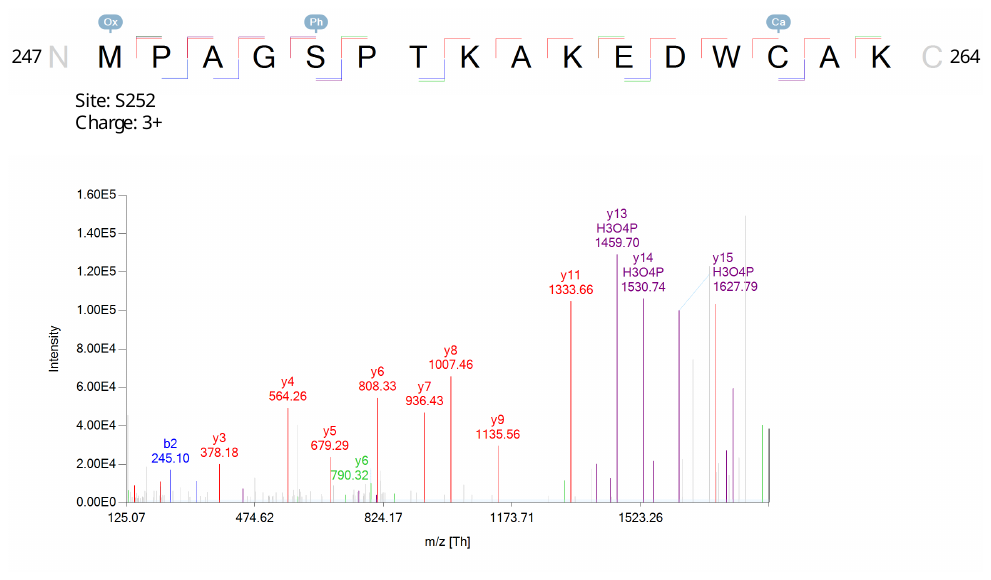

**B**

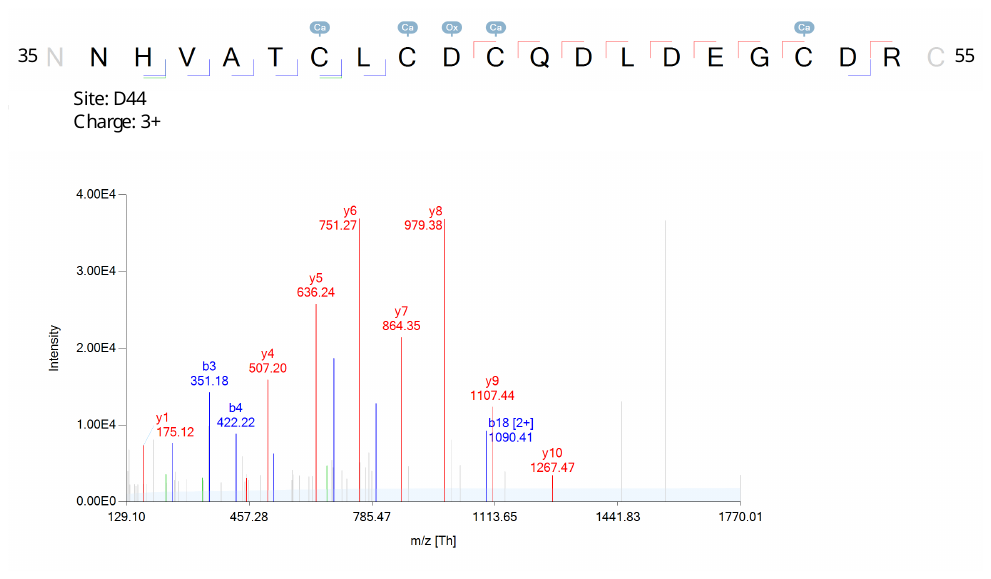

**C**

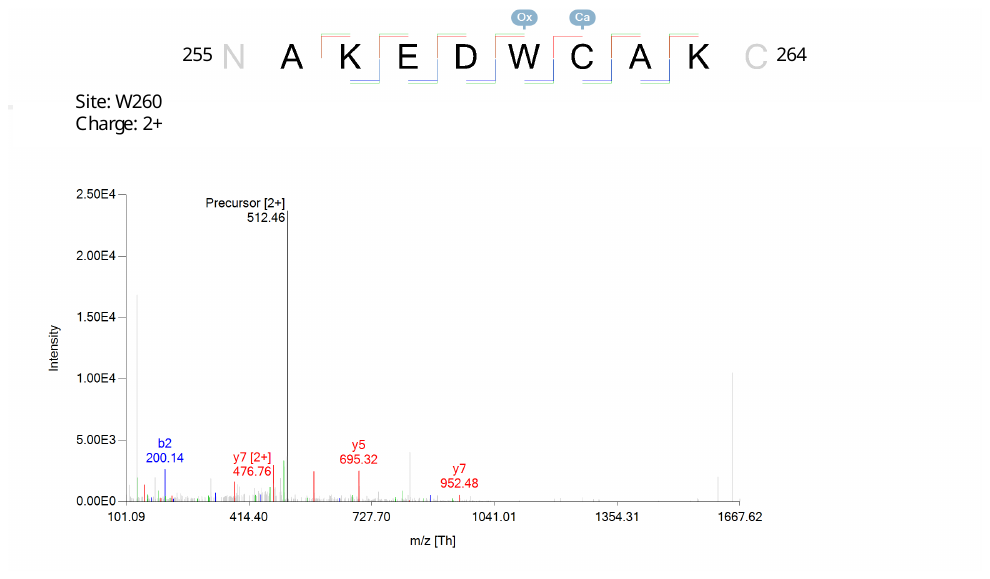

**D**

**Supplementary Figure 2. Annotated mass spectra of zDHHC23 tryptic modified peptides** Annotated deconvoluted tandem mass spectra of tryptic peptide ions from zDHHC23 containing sites of post-translational modification. Assigned HCD product ions are labelled. Peptide sequences and residue numbers are indicated, as are the sites of modification and the isolated peptide ion charge state. Ox = Oxidation, Ph = Phosphorylation and Ca = Carbamidomethylation. **A** Phosphorylation at Ser206 on peptide SLpSSSQLECLSR (Proteome Discoverer ptmRS score = 100). **B** Phosphorylation at Ser252 on peptide MPAGpSPTKAKEDWCAK (Proteome Discoverer ptmRS score = 99.8). **C** Oxidation at Asp44 on peptide NHVATCLCoxDCQDLDEGEDR (PEAKS A Score = 38). **D** Oxidation at Trp260 on peptide AKEDoxWCAK (PEAKS A Score = 1000).
